# Genetic risk of cholangiocarcinoma is linked to the autophagy gene *ATG7*

**DOI:** 10.1101/836767

**Authors:** Stephanie U. Greer, Margret H. Ogmundsdottir, Jiamin Chen, Billy T. Lau, Richard Glenn C. Delacruz, Imelda T. Sandoval, Sigrun Kristjansdottir, David A. Jones, Derrick S. Haslem, Robin Romero, Gail Fulde, John M. Bell, Jon G. Jonasson, Eirikur Steingrimsson, Hanlee P. Ji, Lincoln D. Nadauld

## Abstract

Cholangiocarcinoma is an aggressive cancer originating from the bile duct. Although cholangiocarcinoma does occur in families, to date no specific causative gene has been identified. We identified *ATG7* as a cancer susceptibility gene using a joint genetic analysis of an extended pedigree with familial cholangiocarcinoma in combination with a population genetic association study. Affected family members had a germline mutation (c.2000C>T [p.Arg659*]) in the autophagy related gene, *ATG7*, and all of the affected individuals had cholangiocarcinoma tumors harboring somatic genomic deletions of *ATG7*. From a population genetic study, we identified a germline polymorphism of *ATG7* (c.1591C>G [p.Asp522Glu]) associated with increased risk of cholangiocarcinoma. The autophagy substrate p62 demonstrated a higher accumulation in tumors of p.Asp522Glu carriers compared with non-carriers indicating defective autophagy. To determine whether the germline *ATG7* mutation had functional consequences, we developed an *ATG7*-deficient cholangiocyte cell line, derived from human bile duct, to test for autophagy-mediated lipidation activity. The germline mutation from the familial cholangiocarcinoma demonstrated a lack of lipidation activity compared to the wildtype *ATG7*. Moreover, in zebrafish embryos depleted of *atg7*, a reproducible necrotic head phenotype was rescued by injection of wildtype *ATG7* but not mutant *ATG7*. Our findings point to *ATG7* as a causative genetic risk factor for cholangiocarcinoma and implicate autophagy as a novel cancer driver mechanism.

## INTRODUCTION

Cholangiocarcinoma (CCA; MIM: 615619) is an aggressive, difficult-to-treat epithelial carcinoma arising from the bile duct. Once diagnosed, this biliary duct cancer has a dismal five-year survival rate of less than 10%.^1; 2^ Cholangiocarcinomas are categorized into intrahepatic, perihilar and distal subtypes based on their anatomical location. Perihilar CCA is also referred to as a Klatskin tumor and is anatomically located at the merging of the right and left hepatic bile ducts, and accounts for approximately 50% of all cases.^3^ Genomic studies of cholangiocarcinoma have implicated a variety of known cancer drivers such as *TP53*, *KRAS, ERBB2* and others that play a role in the oncogenic process.^4-7^

There are a number of environmental and genetic risk factors for developing cholangiocarcinoma. Environmental factors, such as chemical exposure or parasitic infections, as well as comorbid conditions, such as primary biliary cholangitis, lead to an elevated risk of CCA.^8; 9^ Regarding genetic risk factors, there are a number of case reports in which known cancer susceptibility syndromes are associated with CCA. There are families in which affected individuals with CCA have germline mutations either in the DNA mismatch repair genes (*MLH1* [MIM: 120436], *MSH2* [MIM: 609309], *MSH6* [MIM: 600678], *PMS1* [MIM: 600258], and *PMS2* [MIM: 600259])^10^ as part of Lynch syndrome (MIM: 120435) or in *BAP1* [MIM: 603089] which is associated with its own distinct tumor predisposition syndrome (MIM: 614327).^11; 12^ However, the majority of families with cholangiocarcinoma remain unexplained with no specific genetic risk factor nor have genomic studies implicated a candidate gene that accounts for genetic risk.^6^

For this study, we identified a family in which three of eight siblings were diagnosed with cholangiocarcinoma with pathologic features consistent with the Klatskin tumor type. Importantly, the affected individuals did not have specific environmental exposures or comorbid conditions associated with increased CCA risk. Given the familial incidence of cholangiocarcinoma in this pedigree, we investigated whether there was a causative germline mutation. To confirm our findings, we conducted a genetic association study of the Icelandic population to identify candidate polymorphisms associated with increased risk for cholangiocarcinoma.

From this combined analysis of familial cholangiocarcinoma and the genetic association study, we identified a germline mutation and specific polymorphism in the *ATG7* gene that are linked to an increased risk of CCA. We used several experimental systems to validate our findings in terms of the functional consequences of these variants. The ATG7 protein plays an essential role in the process of autophagy.^13; 14^ Several core autophagy proteins, including ATG7, are required for an underlying lipidation function that modifies target proteins necessary for autophagy. Studies have implicated that autophagy is protective against tumor initiation, as hepatocyte-specific knockout of the essential autophagy genes results in adenoma formation in the liver.^15; 16^ Our protein and functional studies demonstrated that these variants had an impact on autophagy target proteins. Overall, our results indicate that ATG7 and the autophagy pathway play an important role in the formation of CCA.

## MATERIAL AND METHODS

### Pedigree samples

Our familial genetic study was conducted in compliance with the Helsinki Declaration, and the Institutional Review Board at Stanford University School of Medicine approved the study protocol (76274). Samples were obtained with informed consent. Blood samples were obtained from eight siblings. We obtained matched tumor samples from the individuals affected with CCA (**Table S1** and **Figure 1)**. Genomic DNA extraction was performed using the Promega (Madison, WI) Maxwell 16 Blood DNA Purification Kit and 16 FFPE Plus LEV DNA Purification Kit for the germline blood samples and archival tumor samples, respectively.

**Figure 1.**
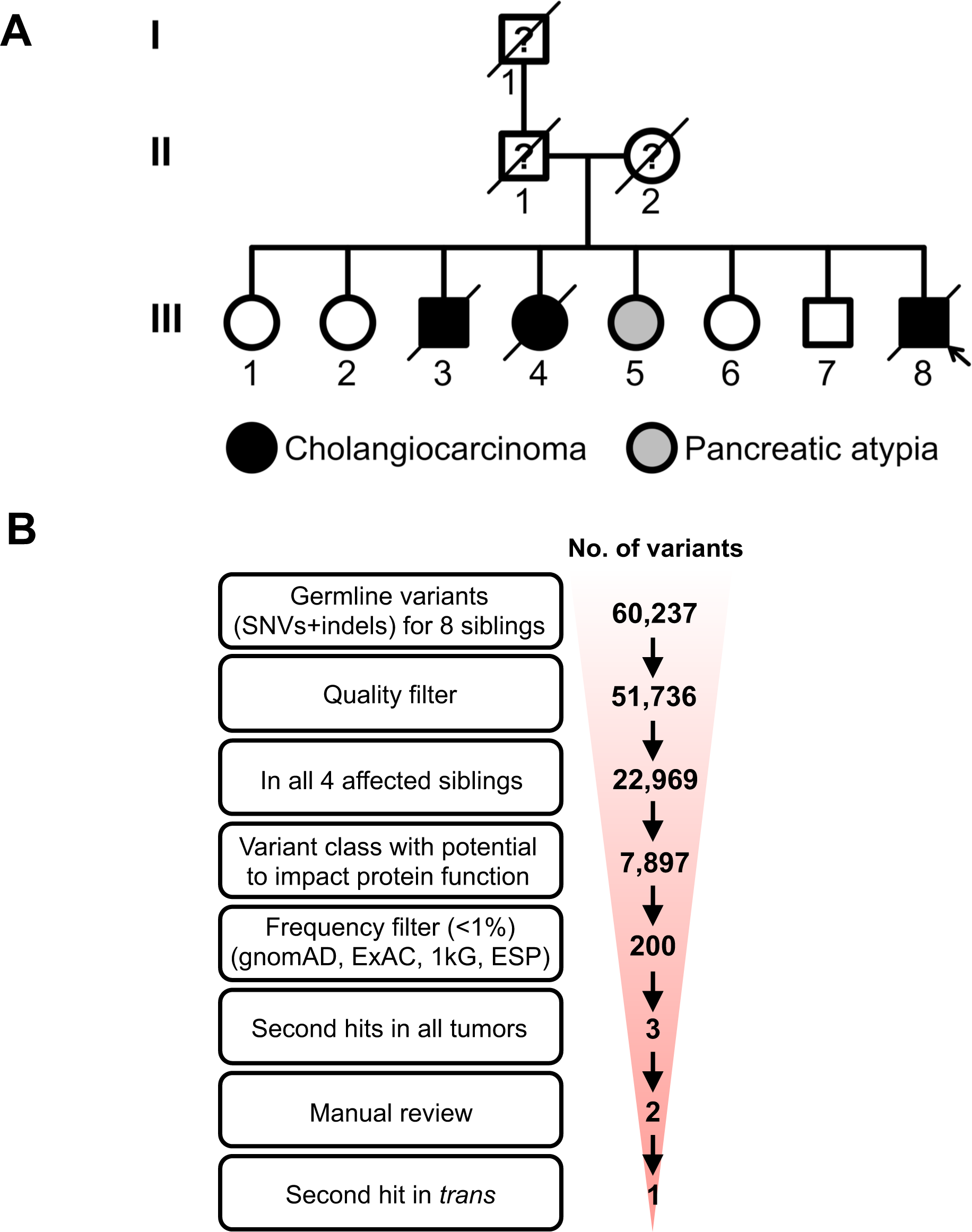
Identification of a putative germline predisposition variant in a family with inherited CCA. **(A) Family pedigree.** This pedigree depicts a family with an inherited predisposition to biliary cancers. The current study had access to samples from generation III of the family. **(B) Candidate filtering.** The number of putative germline variants (single nucleotide variants and insertion-deletions) is indicated for each stage of filtering.

**Table 1.** Association analysis of rs146589465-G

| A1<sup>a</sup> | A2<sup>a</sup> | Freq.<br>A1<sup>a</sup> [%] | Phenotype | N cases | N controls | P-value | OR<sup>a</sup><br>(A1) | 95% CI |
| --- | --- | --- | --- | --- | --- | --- | --- | --- |
| G | C | 0.12 | Cholangiocarcinoma | 353 | 233169 | 1.3×10<sup>-3</sup> | 6.56 | (2.085-20.64) |
|  |  |  | Pituitary adenoma | 422 | 303642 | 3.9×10<sup>-3</sup> | 5.22 | (1.699-16.03) |
|  |  |  | Hepatocellular carcinoma | 280 | 301578 | 2.9×10<sup>-2</sup> | 4.95 | (1.178-20.80) |

Shown are associations for rs146589465-G in the Icelandic cohort that reached a threshold of *P*-value less than 5.0×10<sup>-2</sup>. The variant results in coding change p.Asp522Glu in ATG7.
<sup>a</sup>A1 and A2 stand for the two alleles tested for the marker. Odds-ratio (OR) and population allele frequency (Freq. A1) are provided for allele A1.

### Exome sequencing and germline variant calling

We prepared libraries with the KAPA Hyper Prep kit (Kapa Biosystems, Wilmington, MA) and exome capture was performed with the Nextera Rapid Capture Exome kit (Illumina, San Diego, CA). We sequenced the libraries on either the Illumina HiSeq 2500 system with 100 by 100-bp paired-end reads or the Illumina NextSeq system with 150 by 150-bp paired-end reads. The resulting sequence reads were aligned to human genome build GRCh37.1 with the BWA-MEM algorithm of the Burrows-Wheeler Aligner **(BWA)** v0.7.4.^17^ WES and alignment metrics are available in **Table S2**.

**Table 2.** Analysis of IHC results

| Marker | Nuc/Cytopl | Sample group | 0 | + | ++ | +++ | Total | *P*-value<sup>a</sup> |
| --- | --- | --- | --- | --- | --- | --- | --- | --- |
| ATG7 | Nuclear | Carrier | 4 | 1 | 1 | 0 | 6 | 0.584 |
|  |  | Control | 16 | 5 | 1 | 0 | 22 |  |
|  | Cytoplasmic | Carrier | 0 | 0 | 3 | 3 | 6 | 0.628 |
|  |  | Control | 0 | 3 | 9 | 10 | 22 |  |
| p62 | Nuclear | Carrier | 4 | 0 | 0 | 2 | 6 | 0.638 |
|  |  | Control | 12 | 3 | 2 | 5 | 22 |  |
|  | Cytoplasmic | Carrier | 0 | 1 | 0 | 5 | 6 | 0.003** |
|  |  | Control | 5 | 7 | 8 | 2 | 22 |  |
| LC3B | Nuclear | Carrier | 2 | 3 | 1 | 0 | 6 | 0.638 |
|  |  | Control | 12 | 8 | 2 | 0 | 22 |  |
|  | Cytoplasmic | Carrier | 0 | 2 | 2 | 2 | 6 | 0.590 |
|  |  | Control | 3 | 7 | 3 | 9 | 22 |  |

<sup>a</sup>Chi-square test was performed to assess whether staining patterns differed significantly between carrier and control samples.

We used the Sentieon variant caller^18^ (Mountain View, CA) v201711 to implement the GATK tools, following the GATK Best Practices for variant detection.^19^ We generated a genome variant call format **(gVCF)** file for each sample using Sentieon Haplotyper and then performed joint variant calling on all eight samples with Sentieon GVCFtyper. We followed the GATK best practice recommendations for detecting single nucleotide variants **(SNVs)** and insertion/deletions **(indels)**, and subsequently removed variants without a ‘PASS’ filter. The remaining variants were annotated with the following: Variant Effect Predictor **(VEP)** v74^20^ to assign general information including gene, amino acid, and mutation class; CADD v1.3^21^ to generate a pathogenicity score for each variant; and ESP, ExAC and gnomAD^22^ to determine the population allele frequency of each variant.

To identify germline mutations associated with the affected family members, we used the following criteria: the germline variant was present in all of the affected siblings albeit it could be present in unaffected siblings as well, assuming incomplete penetrance; the variant had the potential to alter protein function (i.e. the variant class had to be one of: missense, nonsense, splice site, stop lost, frameshift, in-frame); and the variant had to occur at low frequency (i.e. less than 1%) in the ‘healthy’ population. This germline filtering strategy is similar to those used in other WES studies to identify putative causal variants in Mendelian disorders.^23^

### Whole genome sequencing with haplotypes

To determine haplotypes over extended genomic segments (i.e. Megabase **(Mb)**) we used linked read whole genome sequencing. This sequencing approach enables one to use barcoded short reads to obtain long-range genomic information from high molecular weight DNA.^24^ With this long-range data, we performed germline structural variant **(SV)** detection and characterized the extended haplotypes across the entire genome for all eight siblings. For all of the pedigree germline samples, we prepared sequencing libraries with the Chromium Gel Bead and Library Kit and the Chromium Instrument (10X Genomics, Pleasanton, CA). These libraries were sequenced on the Illumina HiSeq X system with 150 by 150-bp paired-end reads. The resulting BCL files were converted to fastq files using Long Ranger (v2.1.2) ‘mkfastq’, then Long Ranger ‘wgs’ was run to align the reads to GRCh37.1, detect and phase SNVs/indels, and detect SVs. Linked read sequencing and alignment metrics are available in **Table S2**, and phasing-haplotype metrics are available in **Table S3**.

### Cancer genome sequencing

For the detection of somatic SNVs and indels, we performed high depth WES on three CCA samples in combination with the results from the matched normal blood samples in the germline analysis (above). Sequencing libraries for two of the tumor samples (patient III:3 and III:8) were generated using the KAPA Hyper Prep kit. The other tumor sample (patient III:4) was low quality FFPE, and was prepared using the GemCode Library Kit (10X Genomics) to retain long-range information for improved read alignment. Exome capture was performed with the Nextera Rapid Capture Exome kit (Illumina). We sequenced the libraries on the Illumina HiSeq system with 100 by 100-bp paired-end reads or the Illumina NextSeq system with 150 by 150-bp paired-end reads. The sequence reads were aligned to human genome build GRCh37.1 with the BWA-MEM algorithm of BWA v0.7.4.^17^ For the GemCode library, the BCL files were converted to fastq files using Long Ranger (v2.1.2) ‘mkfastq’, then Long Ranger (v2.1.2) ‘wgs’ was run to align the reads to GRCh37.1.

The mean on-target coverage for the tumor WES data ranged from 88X to 178X; sequencing and alignment metrics are available in **Table S2**. We used the Sentieon v201711 TNsnv tool (similar to the MuTect algorithm) and TNhaplotyper tool (similar to the MuTect2 algorithm) to detect somatic SNVs and small indels using the WES data of each tumor/normal pair, and retained only those variants with a ‘PASS’ filter. We annotated the variants with the same methodology as in the ‘Germline analysis’ (above), and also applied the same filters: the variant had to have the potential to affect protein function and the variant had to occur at low frequency (i.e. less than 1%) in population frequency data. For patient III:4, we used the WES data to identify CNVs using the CNVkit tool.^25^ We annotated the CNV output to determine which genes may have experienced copy loss due to a deletion.

We performed WGS on tumor-normal pairs from patients III:3 and III:8. Sequencing libraries were generated using the KAPA Hyper Prep kit (Kapa Biosystems). The libraries for patient III:3 were sequenced on the Illumina HiSeq 4000 system with 150 by 150-bp paired-end reads and the libraries for patient III:8 were sequenced on the Illumina HiSeq X system with 150 by 150-bp paired-end reads. The resulting sequence reads were aligned to human genome build GRCh37.1 with the BWA-MEM algorithm of BWA v0.7.4.^17^ WGS and alignment metrics are available in **Table S2**. To identify candidate genes where the second (i.e. ‘wildtype’) allele could have been lost due a genomic deletion or gene conversion, we ran the BICseq2^26^ CNV detection tool on these tumor-normal pairs.

We used linked read WGS to retain long-range genomic information from one of the tumor samples (patient III:8). We prepared the sequencing library for the tumor sample using the Chromium Library Kit (10X Genomics). The library was sequenced on the Illumina HiSeq X system with 150 by 150-bp paired-end reads. The resulting BCL files were converted to fastq files using Long Ranger (v2.1.2) ‘mkfastq’, then Long Ranger (v2.1.2) ‘wgs’ was run to align the reads to GRCh37.1, detect and phase SNVs/indels, and detect SVs. The sample was sequenced to a depth of approximately 30X; linked read WGS and alignment metrics are available in **Table S2**, and phasing metrics are available in **Table S3**.

We determined whether each candidate causal mutation was located in *cis* or in *trans* with somatic deletion events in the tumor. For this, we used linked read data to generate digital karyotypes as described in Bell *et al*..^27^ This method generates extended haplotypes from Mb segments, by combining adjacent phase blocks extrapolated from allelic imbalances. The Mb haplotypes are quantitatively determined from the differences in the linked read barcode counts among haplotypes. Using this process, we determined the alleles that belonged to each haplotype in regions with genomic deletions.

### Population genetic study of cholangiocarcinoma

Individuals affected with cholangiocarcinoma were identified through the Icelandic Cancer Registry **(ICR)** and samples were obtained with informed consent. All sample identifiers were encrypted in accordance with the regulations of the Icelandic Data Protection Authority. Approval for the study was granted by the Icelandic National Bioethics Committee (ref. 00/097) and the Icelandic Data Protection Authority (ref. 2001020223).

We identified all of the known variants occurring in *ATG7* for the Icelandic population as previously described.^28^ A total of 31.6 million SNVs and short indels that met quality control criteria were identified in the genomes of 8,453 sequenced Icelanders. These variants were then imputed into 150,656 Icelanders genotyped with Illumina single nucleotide polymorphism **(SNP)** chips, and their genotypes were phased using long-range phasing.^29; 30^ To increase the sample size and power for genetic association analysis, we used genealogical deduction of carrier status for 294,212 relatives lacking array-based genotypes. Subsequently, *ATG7* variant association testing for case-control analysis was performed using logistic regression. The quality of the imputation was evaluated by comparing imputed genotypes to genotypes obtained by direct genotyping. Individual *ATG7* genotyping was performed by applying the Centaurus (Nanogen) platform.

### Immunohistochemistry of ATG7, LC3B and p62

Immunohistochemistry was performed on 3 μm sections from paraffin-embedded tumors. Following deparaffinization, samples were rehydrated and subjected to heat-induced epitope retrieval **(HIER)**. Tris/EDTA buffer pH 9 was used for the ATG7 (clone EP1759Y, Millipore 04-1055; 1:1000), LC3B (clone D11, Cell Signaling Technology 3868; 1:250) and p62 (Enzo PW9860; 1:100) antibodies in a 98.2°C water bath (5/20/20). Endogenous peroxidase activity was blocked with 3% hydrogen peroxide. After incubation with the respective primary antibodies for 30 min at room temperature, EnVision FLEX Kit (DAKO, Glostrup, Denmark) was used for detection. Microscope images of stained sections were scored in a blinded manner. A chi-square test was performed to assess whether staining patterns significantly differed between carrier and control samples.

### Cell culture and DNA/RNA extraction

The immortalized human cholangiocyte cell line MMNK-1, and mouse embryonic fibroblasts **(MEF)** *Atg7^-/-^* and MEF *Atg7^+/+^*, were obtained from RIKEN BioResource Center Cell Bank (Wako, Saitama, Japan). Cells were maintained in DMEM supplemented with 10% fetal bovine serum **(FBS)** at 37°C and 5% CO2. The genomic DNA and total RNA were extracted from cells using the Maxwell 16 Cell LEV DNA Purification Kit and the Maxwell 16 Cell LEV Total RNA Purification Kit (Promega), respectively.

### Generation of isogenic MMNK-1 ATG7^-/-^ cell lines

Using CRISPR-Cas9, an *ATG7* deletion of exon 2 in the MMNK-1 cell line was generated by ZeClinics (Barcelona, Spain). Two sgRNAs targeting exon 2 of *ATG7* were cloned into the PX458 and PX459 vectors. The guide RNA sequences are listed below: *ATG7* sgRNA22_Top: 5’- CACCGAACTGCAGTTTAGAGAGTCC -3’ *ATG7* sgRNA22_Bottom: 5’- AAACGGACTCTCTAAACTGCAGTTC -3’ *ATG7* sgRNA119_Top: 5’- CACCGAAGCTGAACGAGTATCGGC -3’ *ATG7* sgRNA119_Bottom: 5’-AAACGCCGATACTCGTTCAGCTTC -3’

MMNK-1 cells were transfected with both sgRNA vectors and then selected with puromycin. We sorted single MMNK-1 *ATG7^-/-^* cells into 96 wells to screen and isolate isogenic MMNK-1 *ATG7*^-/-^ cells.

### ATG7 expression studies

The plasmid pCMV-myc-Atg7(2) expressing ATG7 isoform 2 was a gift from Toren Finkel (Addgene plasmid #24921).^31^ The plasmid pCMV-myc-Atg7(1) expressing ATG7 isoform 1 was derived from pCMV-myc-Atg7(2) using the Q5 site-directed mutagenesis kit (NEB, Ipswich, MA). Subsequently, we used pCMV-myc-Atg7(1) to introduce the ATG7 mutations p.Arg659*, p.Asp522Glu and p.Cys572Ser using the Q5 site-directed mutagenesis kit. All of the *ATG7* mutations in the vectors were confirmed with Sanger sequencing. The pCMV3-C-GFPSpark vector was used as the control (Sino Biological, Beijing, China). The primers are listed below: *ATG7* p.Arg659*_F: 5’- GTTCTTGATCAATATGAATGAGAAGGATTTAAC -3’ *ATG7* p.Arg659*_R: 5’- TTTGGAAGAACAAGCTGTACATTTG -3’ *ATG7* p.Asp522Glu_F: 5’- AGCTGGGGAGTTGTGTCCA -3’ *ATG7* p.Asp522Glu_R: 5’- CCTTGCTGCTTTGGTTTCTTCA -3’ *ATG7* p.Cys572Ser_F: 5’- CTTGGACCAGCAGTCCACTGTGAGTCG -3’ *ATG7* p.Cys572Ser_R: 5’- GTCCGGTCTCTGGTTGAATCTCCTGG -3’

Human MMNK-1 *ATG7*^-/-^ or mouse MEF *Atg7^-/-^* cells were transfected with 2.5 μg of plasmid expressing WT ATG7 (isoform 1), ATG7 p.Arg659*, ATG7 p.Asp522Glu, ATG7 p.Cys572Ser or GFP control, using Lipofectamine 2000. Cells were starved in EBSS for 3h and harvested for protein extraction 24h post-transfection. Cell lysates (20 μg) were separated on 4–20% precast polyacrylamide gels (Bio-Rad, Hercules, CA) and were transferred to nitrocellulose membranes. The following antibodies were used for immunoblotting: ATG7 (D12B11, Cell Signaling, Danvers, MA), LC3B (D11, Cell Signaling), Myc-Tag (9B11, Cell Signaling), and GAPDH (14C10, Cell Signaling).

Droplet digital PCR **(ddPCR)** was carried out on the Bio-Rad EvaGreen QX200 ddPCR system following the manufacturer instructions. Briefly, each 22µL ddPCR reaction uses 1X EvaGreen ddPCR super-mix, 100nM of primers and 10^-5^ ng of plasmid. The PCR master mix was partitioned and plated per Bio-Rad’s QX200 droplet generation protocol. After thermal cycling, the plate was transferred and read by the Bio-Rad QX200 Droplet Reader.

The expression of mRNA transcripts was determined using cDNA reverse-transcribed from total RNA and quantitative PCR **(qPCR)** was performed on the QuantStudio 6 Flex Real-Time PCR System (Thermo Fisher Scientific, Waltham, MA) using SYBR Green master mixes (Bio-Rad) following the manufacturer instructions. The *RPP30* gene was used as a housekeeping control. PCR products from cDNAs were run and visualized on the Invitrogen E-Gel system (Carlsbad, CA). The primers used for ATG7 ddPCR and cDNA amplification are listed below: ATG7_F: 5’-AAGCCATGATGTCGTCTTCC-3’ ATG7_R: 5’- TCCTTGCTGCTTTGGTTTCT-3’

### *Atg7* knockdown and complementation with human *ATG7* in a zebrafish model

The use of zebrafish in these studies was in accordance with an approved IACUC protocol (#17-03) and within institutional guidelines. Wildtype zebrafish (*TU Danio rerio*) were maintained under standard conditions in accordance with institutional, national ethical, and animal welfare guidelines. Morpholino **(mo)** oligonucleotides were obtained from Gene Tools LLC (Philomath, OR) and solubilized to 1 mM or 3 mM stock solutions in 1x Danieau buffer. Primers were designed according to guidelines recommended by Gene Tools to amplify WT and splice-blocked morphant bands. For rescue experiments, full-length human RNA transcripts for wildtype human *ATG7* as well as p.Cys572Ser and p.Arg659* variants were transcribed from linearized DNA using the mMACHINE transcription kit (Ambion, Austin, TX). One nanoliter of each RNA transcript was injected into 1-2 cell stage embryos. Knockdown of zebrafish *atg7* gene expression and overexpression of mRNA transcripts were assessed by PCR. Zebrafish were scored as ‘rescued’ if they had a head and a set of eyes at 48 hours post-fertilization **(hpf)**, otherwise they were scored as ‘not rescued’.

Automated *in situ* hybridization using the InSituPro Vsi (INTAVIS, Cologne, Germany) was performed using digoxigenin-labeled riboprobes for *atg7*, *dlx2* (*distal-less homeobox 2*), *otx2* (*orthodenticle homeobox 2*), *ascl1a* (*achaete-scute family bHLH transcription factor 1a*), and *fabp10 (fatty acid binding protein 10a, liver basic)*. Embryos were cleared in 2:1 benzyl benzoate/benzyl alcohol solution and documented using an Olympus SZX12/DP71 imaging system (Olympus Corporation, Tokyo, Japan). RNA Reference Sequences deposited in ZFIN (RRID:SCR_002560) were used in designing the riboprobes.

The morpholino sequences were: *atg7* mo (5’-AGCTCGTTCTCCAAACTCACCGTTA-3’); control mo (5’ CCTCTTACCTCAgTTACAATTTATA-3’). The primers used are listed below: ATG7 ORF mRNA F: 5’- AATGGCGGCAGCTACGG -3’ ATG7 ORF mRNA R: 5’- TCAGATGGTCTCATCATCGCTC -3’ ATG7 p.Cys572Ser mutagenesis F: 5’- CTTGGACCAGCAGTcCACTGTGAGTCG -3’ ATG7 p.Cys572Ser mutagenesis R: 5’-GTCCGGTCTCTGGTTGAATCTCCTGG -3’ ATG7 p.Arg659* mutagenesis F: 5’- CTTGATCAATATGAAtGAGAAGGATTTAAC -3’ ATG7 p.Arg659* mutagenesis R: 5’-AACTTTGGAAGAACAAGCTGTACATTTG -3’.

## RESULTS

We identified a family with a high incidence of perihilar CCA (**Table S1** and **Figure 1A)**. Perihilar CCA was present in three of eight siblings (III:3, III:4, III:8) in generation III of the family. Our review of the pathology from the tumors taken at surgical resection and CT imaging studies confirmed this diagnosis among these siblings. One of the eight siblings was diagnosed with pancreatic atypia (III:5). The average age at diagnosis in generation III was approximately 60 years. We enrolled and collected blood from all of the siblings in generation III of the family, both affected and unaffected.

Gastrointestinal **(GI)** cancers affected all three generations of this family. Specifically, the parents and the paternal grandfather were reported to have cancers related to the GI tract. The father (II:1) was reported to have had pancreatic and prostate cancer at the time of death, and the mother (II:2) had pancreatic and cecal tumors. The family also reported that the paternal grandfather (I:1) was diagnosed with stomach cancer. Given the age of these cases, we did not have tumor samples available from these individuals.

### Germline variation of the family members

**Figure S1** outlines our approach to identify candidate genes and germline mutations that may account for the increased incidence of CCA in this family. We conducted both exome and whole genome analysis. First, we examined the germline exome variants of the eight siblings. Overall, we identified on average 33,203 SNVs per sibling and 3,994 indels per sibling (**Table S4**). When we combined the variants of all individuals, there were a total of 46,133 and 5,603 high quality SNVs and indels, respectively, in the germlines of the eight siblings.

Approximately 44% (22,969) of the variants were present in all four of the affected siblings (**Table S5** and **Figure 1B)**. Although we treated the sibling with pancreatic atypia (III:5) as affected, we determined that whether III:5 was treated as affected or unaffected did not affect the final result of our analysis. Of the variants in the affected siblings, 34% (N = 7,897) belonged to a variant class with the potential to impact protein function. For the SNVs, there were 6,610 nonsynonymous variants, 656 splice site variants, and 75 variants affecting stop codons (**Table S6**). For the indels, there were 203 in-frame variants, 191 frameshift variants, and 162 splice site variants (**Table S6**). We cross-validated the variants with the ESP, ExAC and gnomAD databases, which allowed us to eliminate population polymorphisms that were unlikely to be the cause of a rare hereditary condition.^32^ In total, 43 SNVs and 157 indels remained as putative germline mutations, present in 148 genes (**Table S5** and **Figure 1B)**.

Second, we used whole genome sequencing to identify germline deletions, rearrangements or other SVs that may be segregating with the affected individuals. We identified 31 germline rearrangements among all eight siblings (**Table S7**), including 16 deletions (mean size ∼ 71 kb), 10 duplications (mean size ∼ 111 kb), three inversions (mean size ∼ 56 kb), and two SVs of an unknown class (mean size ∼ 46 kb). Fourteen SVs were detected in only one individual while the remaining 17 SVs were detected in more than one individual. Eight SVs were detected in the germline genomes of all four affected individuals, including seven deletions and one duplication. Notably, all of the deletions shared by the affected siblings were also shared with all of the unaffected siblings. Within the segment intervals of the genomic deletions, there were no previously identified tumor suppressors or other cancer-related genes (**Table S7**). As noted, a genomic segment with a duplication was found among all of the affected siblings, however this duplication contained no candidate genes related to a gain-of-function role in cancer. Thus, we determined that there were no germline rearrangements that implicated a susceptibility locus.

We sought to determine whether we could identify any candidate germline mutations in this family that affected genes previously identified as germline predisposition genes in CCA. Familial CCA cases typically occur in the context of cancer predisposition syndromes where the risk of multiple cancer types is increased. Familial cases of CCA have been reported in Lynch syndrome caused by mutations in the DNA mismatch repair genes (*MLH1* [MIM: 120436], *MSH2* [MIM: 609309], *MSH6* [MIM: 600678], *PMS1* [MIM: 600258], and *PMS2* [MIM: 600259])^10^ and in BAP1 tumor predisposition syndrome, where the *BAP1* tumor suppressor gene is mutated.^11; 12^ Our germline analysis of this family did not identify mutations in any of these previously reported CCA predisposition genes.

Genetic studies of hereditary pancreatic cancers have included CCA given the similarities of their embryonic tissue origins and epithelial features which involve secretory physiologic functions.^4; 33^ Thus, we determined whether there were any candidate germline mutations in this family in genes previously identified as germline predisposition genes in pancreatic cancer. An estimated 10% of all pancreatic cancer cases are hereditary, frequently as part of known familial cancer syndromes.^34; 35^ Inherited mutations in *BRCA1* [MIM: 113705]^36^, *BRCA2* [MIM: 600185]^37^, *PALB2* [MIM: 610355]^38^, *ATM* [MIM: 607585]^39^, *STK11* [MIM: 602216]^40^, *CDKN2A* [MIM: 600160]^41^, and *BRIP1* [MIM: 605882]^42^ have been linked to familial pancreatic cancer.

We evaluated all genes involved in hereditary pancreatic cancer mentioned above and also genes reported as risk factors for pancreatic cancer from recent genome-wide association studies, including the genes *TNS3* [MIM: 606825], *NOC2L* [MIM: 610770], *HNF4G* [MIM: 605966], *HNF1B* [MIM: 189907], *GRP* [MIM: 137260], *UHMK1* [MIM: 608849], *AP1G2* [MIM: 603534], *DNTA*, *CHST6* [MIM: 605294], *FGFR3* [MIM: 134934], and *EPHA1* [MIM: 179610].^39;43-45^ Our germline analysis of this family did not identify mutations in any of these previously reported pancreatic cancer predisposition genes.

### Cancer exome and whole genome sequencing

To identify biallelic genetic events in the familial tumor samples, we conducted whole exome and whole genome sequencing of the three tumor-normal sample pairs (i.e. patient III:3, III:4, III:8). With the WES data, we identified somatic cancer mutations and other genetic aberrations (**Tables S8-S11**). With the WGS data, we examined genomic copy number changes as a way of ascertaining whether any deletions overlapped with our candidate germline mutations. The tumor of patient III:3 had five somatic copy number deletions with an average size of 49.8 Mb and containing a total of 3050 genes. In patient III:4, there were 32 somatic deletions with an average size of 34.8 Mb containing 9,620 genes. Patient III:8 had 20 somatically deleted regions with an average size of 24.9 Mb and containing 3,053 genes (**Table S12**).

The principal cancer driver mutations identified in the tumors of the affected individuals have been reported in other CCA genomic studies. Tumors of two patients (III:3 and III:8) harbored a biallelic loss of *ARID1A*, a cancer driver gene associated with the molecular subtype identified as ‘cluster 1’ in a genomic study of 489 CCA tumors.^5^ Also mutated in the patient tumors were other reported CCA driver genes, including *PIK3CA*, *FGFR2*, *BAP1*, *PBRM1*, and *CDKN2A* (**Table S9-S12**).^4^

The tumor mutation burden of these samples ranged from 43 to 79 non-silent somatic mutations per sample (**Table S8**), which is similar to the mutation rate reported in a study of 71 CCA tumor samples, where they found an average of 64 nonsynonymous somatic SNVs per sample.^5^ Thus, the tumor mutation burden of the affected individual tumors displayed no evidence of any hypermutation suggesting deficiency of DNA mismatch repair, or an underlying Lynch syndrome.

### Identification of loss-of-heterozygosity using haplotype analysis

We evaluated whether the candidate genes we identified with germline mutations also harbored somatic alterations in the tumors of the three siblings with cholangiocarcinoma. We had only a germline blood sample for the sibling with pancreatic atypia (III:5) and therefore were unable to perform any somatic analysis for this individual. Of the 148 candidate genes with germline mutations, 14, 57, and 22 genes were determined to have somatic mutations in patient III:3, III:4 and III:8, respectively. We identified three genes (*PP2D1*, *SPSB1* [MIM: 611657], and *ATG7*) with both germline mutations and somatic alterations that were identified among all three affected individuals’ tumors (**Table S13-S14**).

We identified a germline missense mutation in the ***PP2D1*** gene (c.1898G>A [p.Pro547Leu]) but it was eliminated as a candidate due to a lack of functional evidence based on variant impact predictions. The *PP2D1* substitution was scored as being benign and tolerant as per PolyPhen and SIFT analysis, respectively, and also had a low CADD score (5.309) denoting low predicted pathogenicity (**Table S14**). With elimination of this gene, two germline candidates remained: a missense mutation in ***SPSB1*** (c.945C>T [p.Arg202Trp]) that occurred in all eight siblings and a nonsense mutation in ***ATG7*** that was present in the four affected siblings as well as two unaffected siblings (III:2 and III:6). For both of these candidate germline variants, all three patient tumor samples harbored somatic copy number deletions across the genic regions (**Table S12-S13**).

Next, we determined the extended haplotypes of each sibling across the genomic regions harboring each of the candidate genes (*SPSB1* and *ATG7*), and used this information to determine which haplotypes were affected by the somatic copy number deletions. As described below, our haplotype results provided supportive evidence that implicated *ATG7* as the only candidate gene demonstrating biallelic events that affected both gene copies (i.e. a germline mutation in one copy and a somatic deletion of the other copy, resulting in loss of heterozygosity).

For the haplotype analysis, we used linked read WGS. As we have previously published, this method uses molecular barcodes to label high molecular weight DNA molecules and subsequently, these barcodes provide indexed sequence reads and thus retain long-range genomic information from those individual high molecular weight DNA molecules.^24^ Using linked read WGS, Mb-scale haplotypes are generated which can be extended to span entire chromosome arms, as we have shown previously.^27^ We identified the haplotypes at the genomic loci for approximately 1 Mb surrounding each candidate gene of interest (*SPSB1* and *ATG7*). We performed pairwise comparisons of the variant allelic content of each haplotype for all eight of the siblings. Thus, we verified that the haplotype containing the *SPSB1* germline mutation was present among all siblings and the haplotype containing the *ATG7* germline mutation was common to the six siblings with the *ATG7* germline mutation (**Figure 2A**).

**Figure 2.**
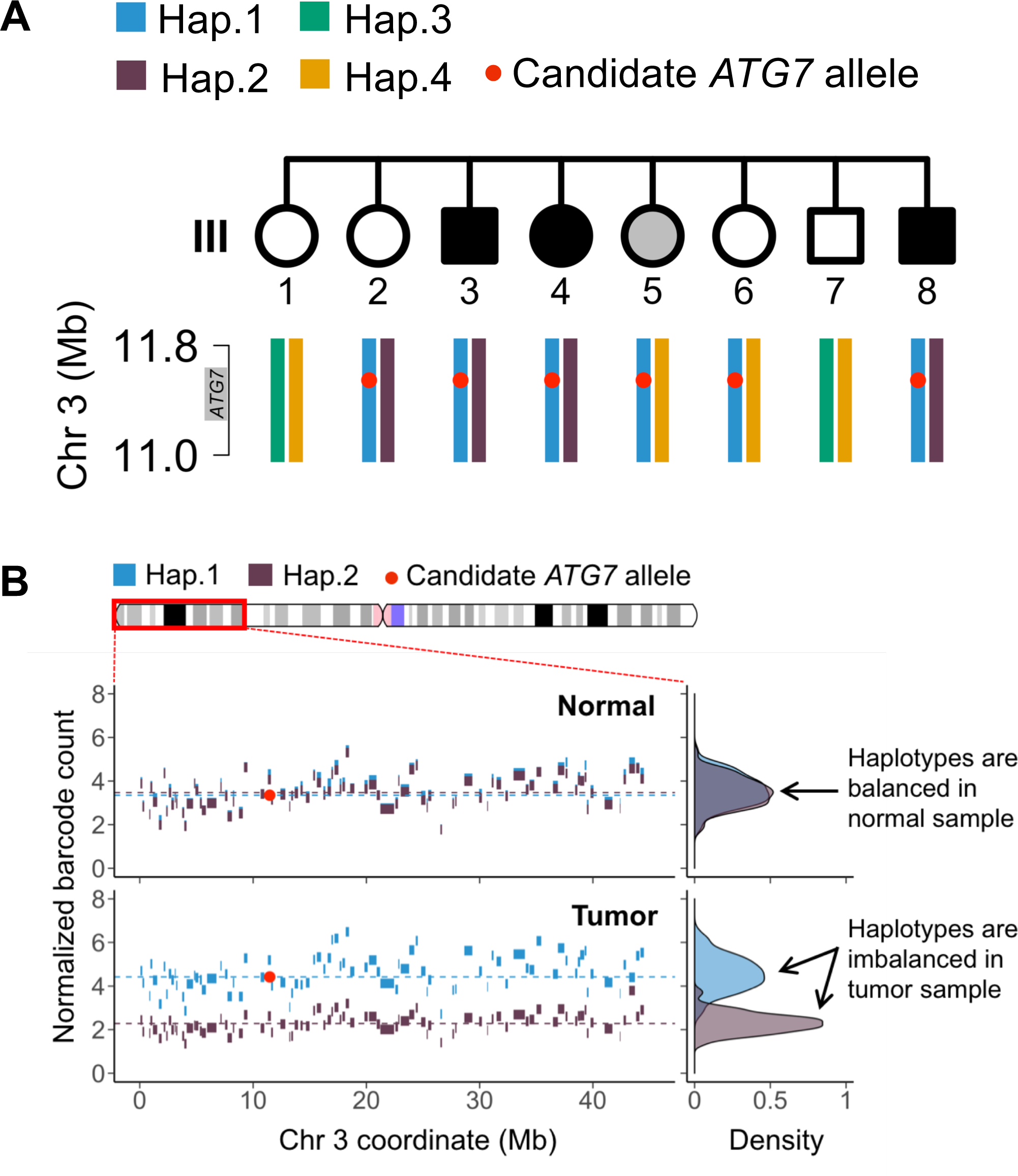
Haplotype analysis of the candidate *ATG7* allele. **(A) Germline haplotype analysis of siblings for the *ATG7* genomic region.** The haplotype segregation across all eight siblings in generation III was determined for the 0.8 Mb genomic region surrounding the *ATG7* allele (red dot). The candidate *ATG7* allele exists in haplotype 1 (blue). **(B) Extended haplotype of the 45 Mb deleted region of chromosome 3p in individual III:8.** The blocks indicate the original fragmented haplotypes, and their color denotes their subsequent assignment to haplotypes covering many Mb. The candidate *ATG7* allele exists in haplotype 1 (blue), which was the non-deleted haplotype in the tumor of this individual.

Next, for the two candidate germline mutations in *SPSB1* and *ATG7*, we determined whether the haplotype of each mutation was in *cis* or in *trans* with the genomic deletion identified in a tumor sample. For this analysis, we used linked read WGS data for the tumor of patient III:8 to ascertain whether the somatic copy number deletions were deletions of the mutant allele or deletions of the wildtype allele. We applied the chromosomal haplotyping method described in Bell *et al*. (2017) where we use somatic allelic imbalances to link phase blocks and identify larger haplotypes.^27^ Our analysis identified 14 chromosome arms where we could determine extended haplotypes covering anywhere from 1% to 98% of a given chromosome arm (**Table S15**). *SPSB1* was located on chromosome arm 1p for which we had an encompassing 36.6 Mb haplotype (p=5.3×10^-13^). *ATG7* was located on chromosome arm 3p for which we had an encompassing 44.4 Mb haplotype (p=2.2×10^-16^)(**Table S15**). To visualize the haplotype structure across these genomic deletions, we plotted the two haplotypes such that individual phase blocks are color-coded according to their assignment to an extended genomic haplotype (**Figures 2B** and **S2**).

In the case of *SPSB1*, we discovered that the germline mutation (p.Arg202Trp) was deleted in the tumor genome and thus the remaining allele was wildtype (**Table S16** and **Figure S2**). Based on this result, we concluded that there was no evidence implicating the *SPSB1* gene as the causal tumor suppressor gene in this family.

Conversely, in the case of ATG7, we found that the wildtype *ATG7* allele was deleted in the tumor and thus the haplotype with the *ATG7* germline mutation (p.Arg659*) was retained (**Table S16** and **Figure 2B**). Together, these two events fulfilled the two-hit criteria of *ATG7* being a putative tumor suppressor gene. Given this result, from our genome-wide survey the *ATG7* germline mutation was our highest-ranking candidate as a potential susceptibility gene in this family.

### Identification of an *ATG7* risk allele for cholangiocarcinoma in the Icelandic population

We leveraged the Icelandic population for a study of *ATG7* association with cholangiocarcinoma. This population is ideal for genetic association studies due to its well-documented genealogy spanning centuries, a genetically homogenous population and the availability of nationwide healthcare records.^46^ We analyzed sequence variants in *ATG7* identified in whole genome sequence data from 8,453 Icelanders sequenced to a median depth of 32x.^28^ Based on the SNVs and SNPs that were identified from the sequencing data, we imputed the *ATG7* genotypes of an additional 150,656 Icelanders with SNP array information. Additional familial imputation, using the nationwide Icelandic genealogical database, allowed these genotypes to be propagated into 294,212 un-genotyped close relatives of the chip-genotyped individuals. Thus, the overall sample size was adequately powered to detect the association of specific *ATG7* variants and polymorphisms with an increased risk of developing CCA. Six coding variants in *ATG7* reached our threshold of imputation quality (**Table S17**). Using information from the ICR, we tested the association between these variants and affected individuals with CCA (*N*=353). This analysis revealed an association between the minor allele of rs146589465 (rs146589465-G) and increased risk of CCA (OR 6.56, *P*=1.3×10^-3^; **Table 1**). We assessed the quality of the imputation of rs146589465 by directly genotyping imputed carriers (*N*=154) and non-carriers (*N*=1180). The concordance between imputed and directly measured genotypes was 0.999 (**Table S18**).

The variant rs146589465-G results in an aspartic acid to glutamic acid substitution at position 522 in the ATG7 protein (p.Asp522Glu). The variant is rare and is present in 1/400 individuals in the Icelandic population (allelic frequency is 0.12%). It is also present at a similar frequency in individuals of European ancestry per the gnomAD database (**Table S17**) and is present only at a very low frequency in other populations, with its highest frequency being ∼0.1% in the Latino population. To assess whether rs146589465 was associated with increased risk for other cancer types, we tested 19 additional tumor types using information from the ICR (**Table S19**). The risk of hepatocellular carcinoma (OR=4.95, *P*=2.9×10^-2^; **Table 1**) and pituitary adenoma (*N*=422; OR=5.22, *P*=3.9×10^-3^) was also elevated with rs146589465-G (**Table 1**). When we tested the association between rs146589465 and a large number of quantitative traits, we found an association between rs146589465-G and bile acid levels (*N*=2183; effect 1.25, *P*=1.3×10^-3^). However, this association was not significant after multiple testing correction.

### The p62 protein accumulates in tumors from rs146589465-G carriers

To determine if the ATG7 rs146589465-G (p.Asp522Glu) variant leads to autophagy defects in patient samples, we analyzed CCA as well as hepatocellular tumors from 6 carriers and 22 non-carriers (**Table S20**). There was no difference in the staining of ATG7 when comparing tumors from carriers and non-carriers, suggesting that the variant does not alter the expression of the ATG7 protein (**Table 2** and **Figure 3**). In addition, there was no difference in expression of the autophagy marker LC3 nor in the pattern of LC3 punctae in the tissue samples (**Table 2** and **Figure 3**).

**Figure 3.**
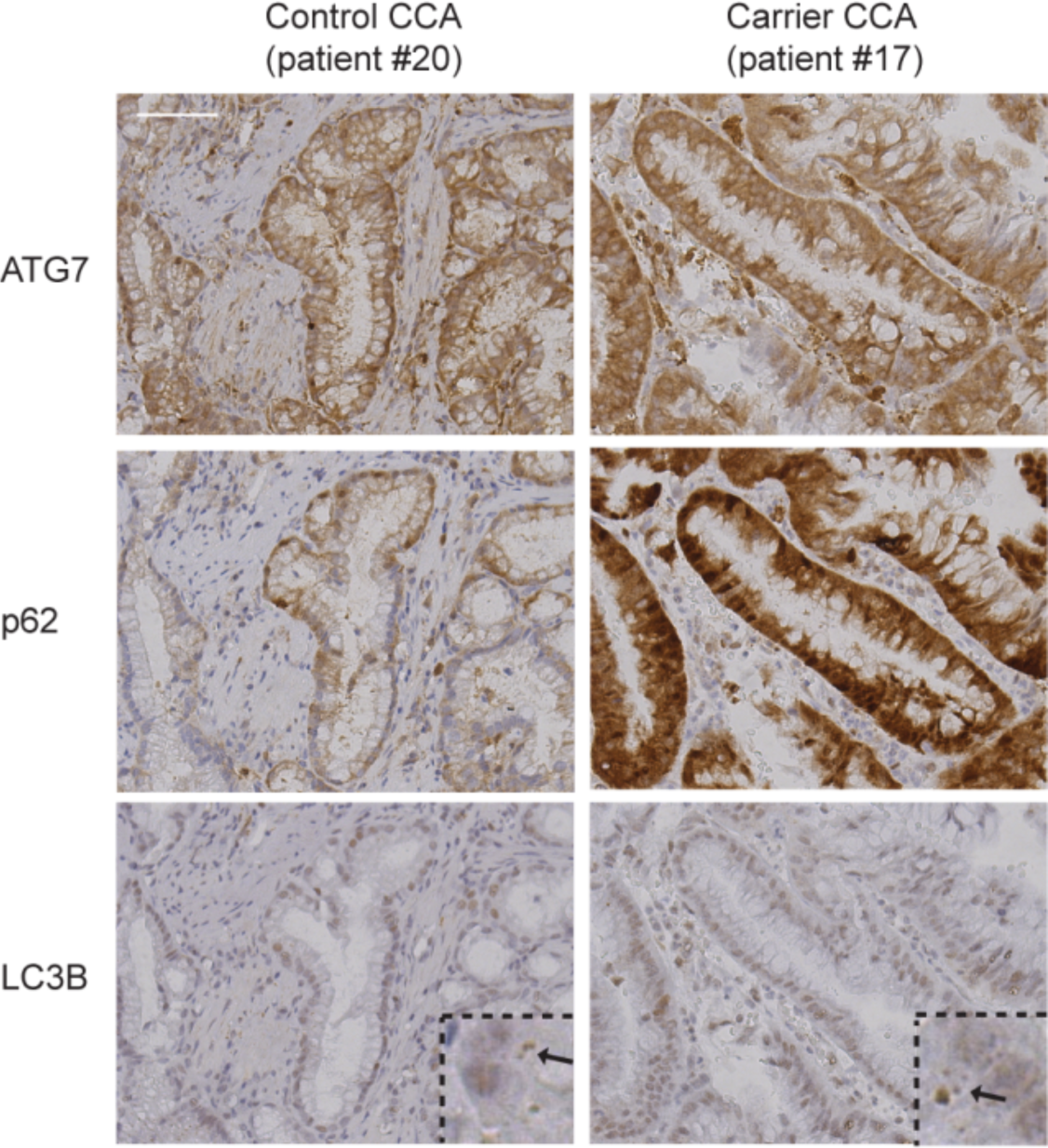
Increased p62 expression in tumors of ATG7 variant carriers. The microscopic images show sections of CCA from an rs146589465 carrier (patient #17) and a non-carrier control (patient #20) stained with antibodies against ATG7, p62 and LC3B. For the LC3B staining, a higher magnification window is shown and the arrows point to LC3 dots that are indicative of autophagosomes. Scale bar represents 20 µm and applies to all panels.

The SQSTM1 protein, also known as p62, serves as an autophagy receptor by linking ubiquitinated cargo to LC3 on the autophagosomal membrane for autophagy degradation.^47^ Since p62 is itself degraded by autophagy, its accumulation is frequently observed as a result of autophagy defects. We conducted immunohistochemistry studies for the presence of p62 in primary tumors among the carriers. We observed significantly higher levels of cytoplasmic p62 protein in samples from rs146589465-G carriers compared to those of non-carriers (*P*=0.003) (**Table 2**, **Figures 3** and **S3**). Accumulation of p62 has also been linked to an antioxidant response in tumor cells and our observation is consistent with increased mouse p62 levels in liver adenomas of liver-specific *Atg7* knockout mice.^48^ To conclude, our findings show that there is greater accumulation of p62 in tumors from rs146589465-G carriers than in non-carriers, indicating an autophagy defect in carriers.

### **The germline ATG7 p.Arg659* mutation leads to a loss-of-function of lipidation activity**

Using SWISS-MODEL^49^, we modeled and annotated the human ATG7 protein structure based on the yeast Atg7 protein (**Figures 4A and S4A**). Human ATG7 has E1-like activity similar to yeast Atg7.^14^ The yeast Atg7 protein has been crystallized, revealing two conserved structural domains, namely the N-terminal domain **(NTD)** and the C-terminal domain **(CTD)** (**Figures 4A and S4A**).^50; 51^ The NTD is responsible for Atg3 binding and the CTD is required for Atg8 recognition and binding. The CTD has two subdomains that include a homodimeric adenylation domain **(AD)** and an extreme C-terminal domain **(ECTD)**. The human ATG7 p.Arg659 residue is located in a highly conserved area of the ECTD, and generates a stop codon leading to truncation of the ECTD (**Figure S4B**). The human ATG7 p.Asp522 residue is located in an exposed exterior loop of the AD, such that p.Asp522Glu may affect protein interactions (**Figure S4C**).

**Figure 4.**
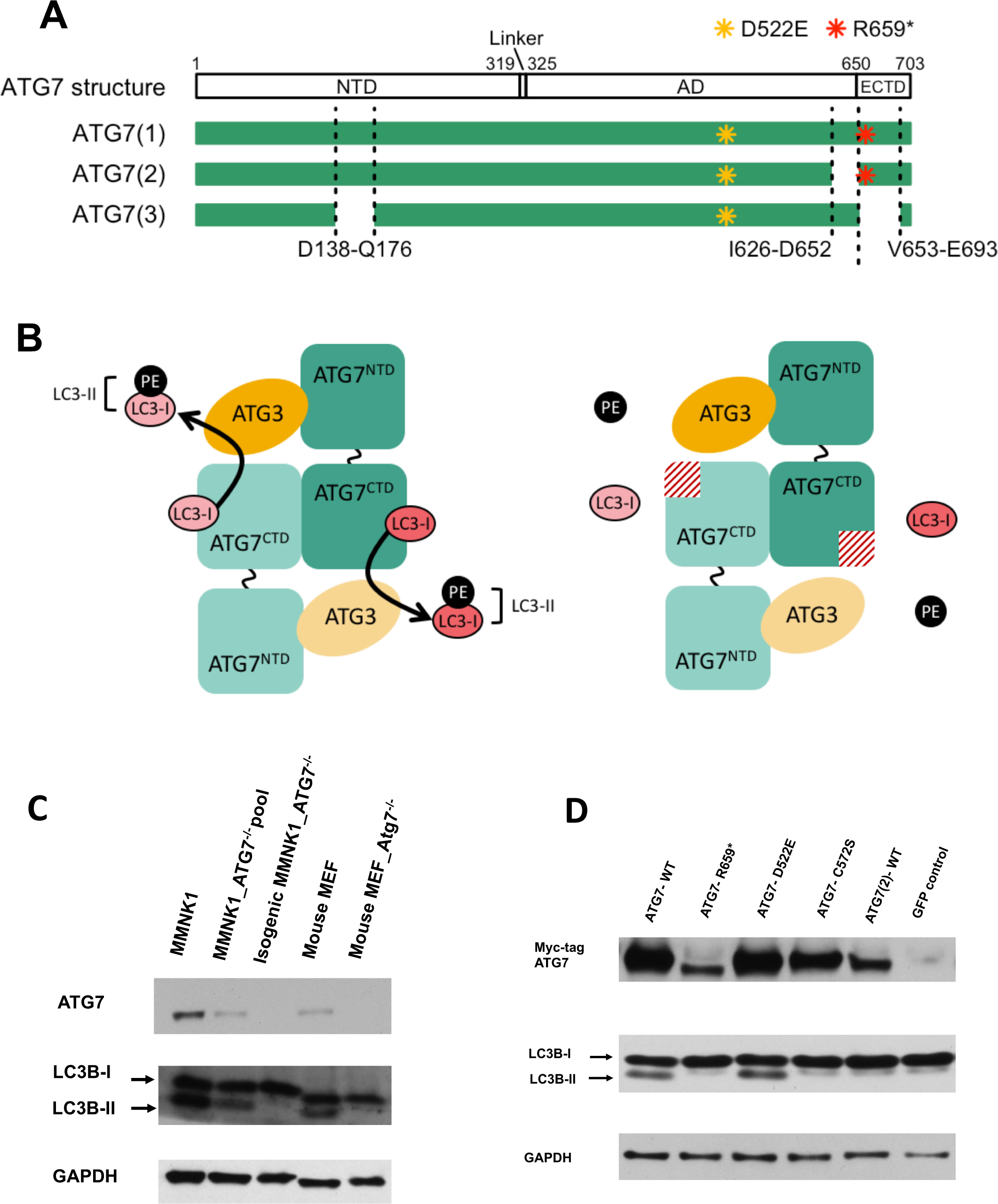
ATG7 p.Arg659* is a loss-of-function mutation. **(A)** The predicted structure of ATG7 contains an N-terminal domain **(NTD)**, linker region, adenylation domain **(AD)**, and extreme C-terminal domain **(ECTD)**. The p.Arg659* (R659*) mutation is present in isoform 1 and isoform 2 of the ATG7 protein, but not in isoform 3. The p.Asp522Glu (D522E) mutation is present in all three isoforms of ATG7. **(B)** Predicted functional impact of ATG7 p.Arg659* mutation. Wildtype ATG7 and ATG3 conjugate ATG8 proteins such as LC3 with phosphatidylethanolamine **(PE)**. We determined that the ATG7 p.Arg659* mutant lacks PE conjugation activity to LC3 as a result of the ECTD truncation. **(C)** MMNK1_ATG7^-/-^ cells did not convert LC3B-I to the lipidated form LC3B-II. Mouse MEF and MEF_Atg7^-/-^ were used as controls. **(D)** Expression of ATG7 WT and p.Asp522Glu restored the lipidation of LC3B whereas expression of ATG7 p.Arg659*, p.Cys572Ser, or wildtype ATG7(2) failed to lipidate LC3B from LC3B-I to LC3B-II. A Myc-tag was used to detect the expression of ATG7 from various vectors. GAPDH was used as the loading control.

Human *ATG7* has three isoforms (**Figure 4A**). Isoforms 1 and 2 are expressed in human tissues whereas isoform 3 is barely expressed.^52^ ATG7 isoform 2 lacks a part of the region required for LC3 binding, and is thus unable to lipidate LC3.^52^ Given that the ECTD is essential for yeast Atg8 recognition and binding^50; 51^, we conducted *in vitro* functional assays to determine whether the ATG7 p.Arg659* variant altered lipidation of LC3 (**Figure 4B**). We also included the ATG7 p.Asp522Glu mutation in our analysis, as well as a known inactivating mutation of ATG7 (p.Cys572Ser).

For our functional assays, we used CRISPR/Cas9 to generate a homozygous deletion in exon 2 of *ATG7* in a human cholangiocyte cell line MMNK-1. This cell line was derived from the human bile duct, thus representing the same primary tissue from which CCA arises. Isogenic MMNK-1 *ATG7^-/-^* cell lines were established by sorting single cells and identifying clones with the targeted deletion by Sanger sequencing (**Figure S5A**) and targeted amplicon sequencing (**Figure S5B**). Furthermore, we confirmed the complete loss of ATG7 protein expression in MMNK-1 *ATG7^-/-^* cells (**Figure 4C**).

LC3B lipidation can be monitored by PAGE and Western blotting since the PE-bound lipidated LC3B-II migrates faster than the unlipidated LC3B-I. As expected, MMNK-1 ATG7^-/-^ cells did not have LC3B lipidation activity, evident by the lack of a lower LC3B-II band (**Figure 4C**). To evaluate which ATG7 variants could rescue lipidation activity, MMNK-1 ATG7^-/-^ cells were transfected with vectors expressing wildtype ATG7 isoform 1, p.Arg659* (the familial variant), p.Cys572Ser (a known inactivating mutation of ATG7), p.Asp522Glu (the Icelandic population variant), and wildtype ATG7 isoform 2. The expression level of p.Arg659* was notably lower than the wildtype ATG7, despite similar plasmid input and mRNA expression (**Figure S6**), suggesting that the truncating mutation may affect the stability of the protein. Expression of either wildtype ATG7 isoform 1 or p.Asp522Glu ATG7 enabled conversion of LC3B-I to LC3B-II in MMNK1 ATG7^-/-^ cells. In comparison, expression of p.Arg659*, p.Cys572Ser, or ATG7 isoform 2 did not enable the lipidation of LC3B-I to LC3B-II (**Figure 4D**). In conclusion, the p.Arg659* candidate familial mutation resulted in a loss-of-function of the ATG7 lipidation activity.

### *ATG7* p.Arg659* failed to rescue *atg7* knockdown in zebrafish

Alignment of the human and zebrafish ATG7 protein sequences shows that the region truncated due to the p.Arg659* mutation is conserved between the two species (**Figure S7**). In contrast, residue p.Asp522 is not conserved. The catalytically active Cys572 residue is conserved. To further demonstrate the functional impact of the *ATG7* p.Arg659* mutation, we tested whether this mutation could rescue the phenotypic effect of *atg7* knockdown in zebrafish embryos. Previous studies of zebrafish have demonstrated its utility as a genetic model system for the process of autophagy as well as for modeling human cancers, including pancreatic cancer.^53; 54^ Zebrafish are an attractive and amenable model system due to their rapid reproduction, which results in hundreds of optically transparent embryos that develop externally.^55^ We first determined that *atg7* is expressed in the head region of the 24 hpf zebrafish embryo (**Figure 5A**). Subsequently, knocking down zebrafish *atg7* gene expression using an antisense oligonucleotide, we observed reproducible head necrosis (**Figure 5B**) and decreased expression of three brain markers (**Figure 5C**). Co-injection of human *ATG7* WT mRNA reduced the number of embryos with the necrotic head phenotype (p=0.0002). In comparison, the cDNA with either the *ATG7* p.Arg659* mutation or the known loss-of-function mutant p.Cys527Ser did not rescue this phenotype (**Figure 5D**). This result showed that *ATG7* p.Arg659* is a loss-of-function mutation *in vivo*. Finally, we verified that *ATG7* plays a role in liver development by performing *in situ* hybridization. Using the liver marker *fabp10*, we observed that 72 hpf *atg7* morphant embryos consistently had smaller livers (100%; n=17) compared to control embryos (100%; n=9) (**Figure 5E**).

**Figure 5.**
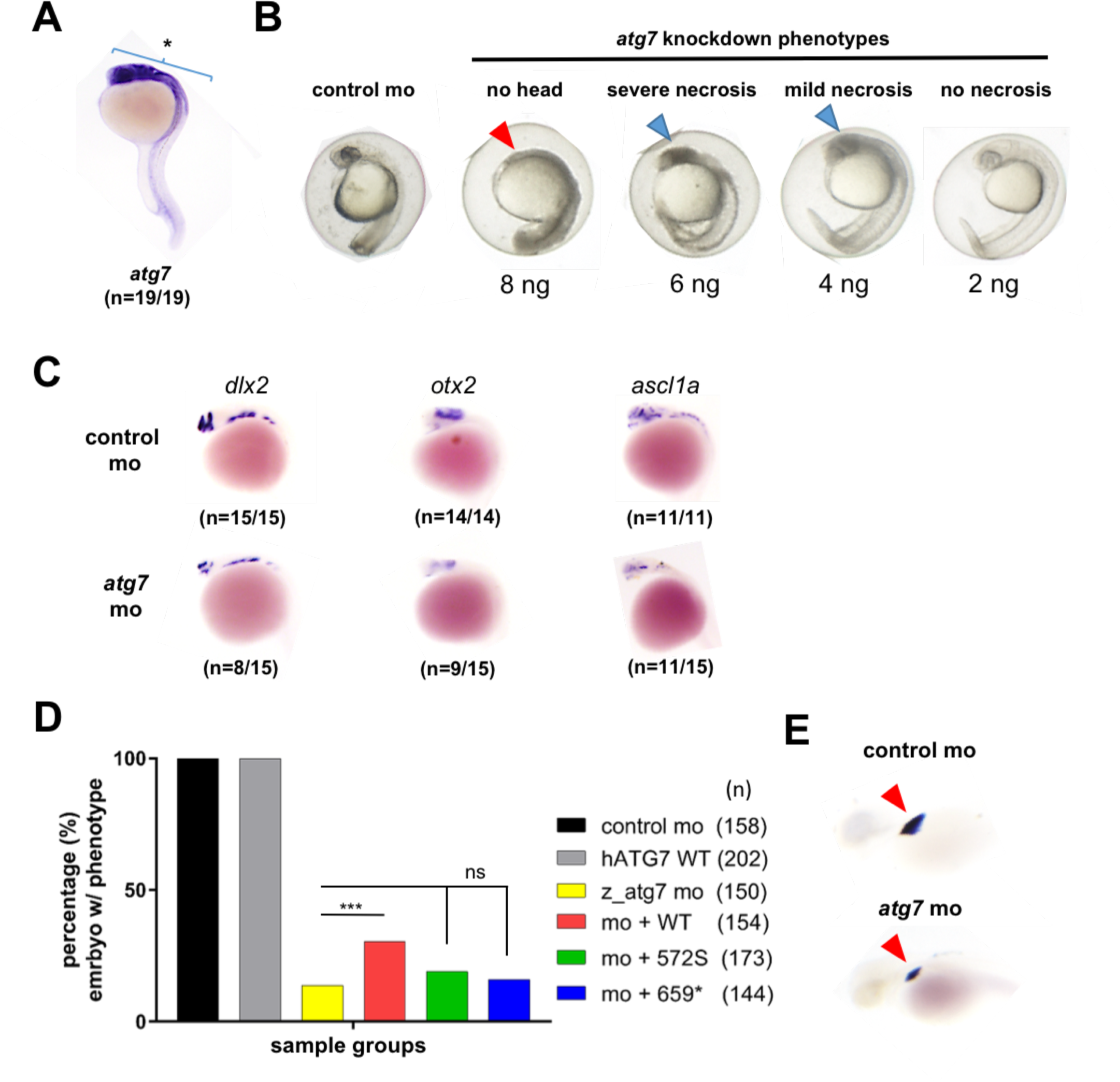
Human wildtype *ATG7* cDNA but not the p.Cys572Ser and p.Arg659* variants complement an *atg7* phenotype in zebrafish. **(A)** Representative *atg7* expression (*) in 24h old wildtype embryos probed by *in situ* hybridization (head region = *). **(B)** Two to eight ng *atg7* mo was injected into 1 - 2 cell stage zebrafish embryos that were then allowed to develop overnight. At 24 hours post-fertilization **(hpf)**, embryos injected with 8 ng *atg7* mo were headless (red arrowhead) while at lower doses (6 and 4 ng), reproducible brain necrosis (blue arrowhead) was observed. **(C)** Representative figures showing decreased staining for three brain markers (*dlx2*, *otx2*, and *ascl1a*) in 24 hpf *atg7* morphants (bottom row) compared to control (top row) by *in situ* hybridization. **(D)** cDNAs expressing WT ATG7, or the p.Cys572Ser or p.Arg659* variants were separately co-injected with 6 ng *atg7* mo. The *ATG7* WT construct rescued the brain necrosis phenotype (***p=0.0002), whereas neither p.Cys572Ser nor p.Arg659* was able to rescue (ns). Fisher’s exact tests were used to determine statistical significance. A p-value less than 0.05 was considered significant. **(E)** *In situ* hybridization using the liver marker *fabp10* shows that atg7 knockdown led to smaller liver (100%; n = 9/9) compared to control (100%; n = 17/17) in 72 hpf zebrafish embryos.

## DISCUSSION

Leveraging an extended pedigree of familial CCA and a genetic association study, our results implicate the *ATG7* gene as a novel genetic risk factor for elevated CCA risk. This represents the first identification of a germline mutation associated with cholangiocarcinoma and subsequently validated in a separate population genetic analysis. We also confirm autophagy pathway perturbation as a novel cancer driver mechanism in human tumorigenesis.

We identified a family pedigree in which multiple siblings were affected with CCA. To discover candidate germline mutations, we conducted an integrated analysis involving whole exome sequencing, whole genome sequencing, and linked read sequencing. The combined dataset enabled us to evaluate both germline mutations and other genomic alterations such as rearrangements. Our analysis eliminated previously identified familial CCA and pancreatic cancer genes as the potential cause of increased risk. Instead, we identified a deleterious germline mutation of *ATG7* that was associated with the affected individuals, including the sibling with pancreatic atypia. Our genomic study of the affected individuals’ tumors provided supporting data for biallelic somatic events that lead to *ATG7* loss-of-function. Using an independent genetic association study in the Icelandic population, an *ATG7* missense polymorphism (p.Asp522Glu) was associated with the development of CCA.

For this study, we demonstrated that linked read WGS is a useful tool for evaluating candidate causal variants. First, linked read sequencing phases variants across large genomic ranges thus generating Mb-scale haplotypes. These large-scale haplotypes can be compared among family members to identify haplotypes shared by all affected family members. In particular, we verified that each candidate causal germline allele was inherited by all affected individuals in the context of a single haplotype, as would be expected in a Mendelian disorder. For both candidate alleles in *ATG7* and *SPSB1*, we were able to verify that they were inherited in a single haplotype. Linked read sequencing also provided a means to determine whether the somatic variants identified in the tumor samples occurred in *cis* or in *trans* with the candidate germline haplotypes, so we could determine whether a somatic variant represented a second hit. By using linked read barcode counts to determine which allele was deleted, we determined that the *SPSB1* somatic deletion was a deletion of the mutant germline allele and thus not a second hit, whereas the *ATG7* deletion represented a true second hit.

The biological importance of *ATG7* is evident by the neonatal lethality of *Atg7* knockout in mice.^15^ Both the *ATG7* p.Arg659* and p.Asp522Glu mutations identified in this study are heterozygous, further emphasizing the importance of the ATG7 protein. Tissue-specific *Atg7* knockout in mouse models have shown that a lack of Atg7 in hepatocytes leads to adenoma formation.^15^ It would be of interest to investigate the phenotype of mice with *Atg7* knockout in cholangiocytes and whether these mice develop adenomas.

The *ATG7* p.Asp522Glu variant resulted in the accumulation of p62 in the tumors of carriers compared with non-carrier tumors. The p62 protein is a known substrate of autophagy and is commonly used as a reporter for autophagic activity, whereby p62 accumulation indicates reduced autophagy activity.^56^ Interestingly, adenoma formation in *Atg7* hepatocyte knockout mice has been linked with accumulation of p62.^57; 58^ Accumulation of p62 results in the activation of antioxidant response, in addition to c-Myc and mTORC1 activation and metabolic reprogramming beneficial to cancer cells.^59; 60^

The E1-like enzymatic activity of ATG7 resulting in the lipidation of LC3 is the best characterized function of the protein and key to autophagy activation. We characterized the functional properties of variant ATG7 proteins in a human bile duct cell line with a deletion in *ATG7*. Our study demonstrated that wildtype ATG7 and ATG7 p.Asp522Glu were able to rescue LC3 lipidation, whereas ATG7 p.Arg659* and another known ATG7 loss-of-function substitution (p.Cys572Ser) were not able to rescue LC3 lipidation. The p.Asp522Glu population variant is a subtle change from aspartic acid to glutamic acid, whereas p.Arg659* results in a truncation of the ATG7 protein with a more severe functional consequence. Although ATG7 p.Asp522Glu functions like the wildtype protein with regard to LC3 lipidation in cell culture, there was an accumulation of p62 in the tumors of p.Asp522Glu carriers compared with control samples. This indicates defective autophagy-related function of ATG7 p.Asp522Glu in individuals carrying the mutation and suggests accumulative effects in cells expressing the subtle variant protein over a long period of time. Interestingly, we found that knockdown of *atg7* in zebrafish embryos yielded a reproducible phenotype that was rescued by wildtype ATG7, but not by p.Arg659* or another known loss-of-function variant of ATG7 (p.Cys572Ser). This emphasizes the deleterious effect of ATG7 p.Arg659* *in vivo*.

In addition to its key role in autophagy, ATG7 has been shown to bind TP53 directly and regulate subsequent transcription of the critical cell-cycle inhibitor CDKN1A, also referred to as p21.^61^ Mouse embryonic stem cells deficient in Atg7 fail to undergo cell cycle arrest, a defect not seen in Atg5- or Atg6-deficient embryos. The ability to directly bind to TP53 and regulate cell cycle control is one potential mechanism by which the absence of ATG7 could lead to the initiation of tumorigenesis. For future studies, we will investigate the precise mechanism of how *ATG7* loss-of-function contributes to tumorigenesis.

## SUPPLEMENTAL DATA

Supplemental Data include 7 figures and 20 tables.

## ACKNOWLEDGMENTS

The authors recognize and appreciate the patients and families who contributed to the current study. HPJ and LDN had full access to all of the data in the study and take responsibility for the integrity of the data and the accuracy of the data analysis. This work was supported by the following grants from the NIH: P01HG000205 to SUG and HPJ; 1U01CA15192001-A1 to HPJ; 1U01CA176299 to HPJ; HG006137-07 to HPJ; R01 CA116468NIH to DAJ. An award from Intermountain Healthcare supported SUG, JC and HPJ. JC and HPJ were also supported by a Research Scholar Grant (RSG-13-297-01-TBG) from the American Cancer Society. HPJ received additional research support from the Clayville Foundation and the Gastric Cancer Foundation. DAJ received additional funding from the Samuel Waxman Cancer Research Foundation, Oklahoma Center for Adult Stem Cell Research (OCASCR) and Oklahoma Medical Research Foundation (OMRF). LDN was supported by NIH/NCI (5K08CA166512), the Conquer Cancer Foundation (Young Investigator Award), the Gastric Cancer Foundation, and the Carl Kawaja Foundation. This work was supported by grants from the Research Fund of Iceland (no 130230-0529 to ES and MHO, no 184861-052 to ES and 184727-051 to MHO). MHO is also supported by a grant from the Icelandic Cancer Society Research Fund. We acknowledge the Icelandic Cancer Registry for assistance in the ascertainment of the Icelandic cancer patients. We thank deCODE genetics for access to data and facilities, assistance with data analysis and helpful discussions.

## Web Resources

Gene Tools, http://www.gene-tools.com/ ICR, http://www.krabbameinsskra.is OMIM, http://www.omim.org/, ZFIN, https://zfin.org

## Accession Numbers

The accession number for the datasets reported in this paper is National Institute of Health’s dbGaP: phs001593.v1.p1

## Author contributions

LDN and DSH, with assistance from RR and GF, collected the patient data and samples for the family with CCA. BTL conducted the whole genome, whole exome and linked read whole genome sequencing assays. SUG and HPJ analyzed the sequencing data and performed the genetic analysis; the mega-haplotyping analysis was performed by JMB. Statistical analysis of Icelandic population data was performed by MHO. Selection and preparation of paraffin samples, staining, imaging and scoring of patient samples was performed by JGJ, SK, and MHO. The Icelandic population study and FFPE staining experiments were designed and results interpreted by MHO and ES. Cell line experiments were designed and performed by JC. Zebrafish experiments were designed and performed by RCD, ITS, and DAJ. SUG, MHO, ES, JC, DAJ, HPJ, and LDN wrote the manuscript. LDN and HPJ designed, supervised, and coordinated all aspects of the research. All authors revised, read, and approved the final manuscript.

---

**Figure S1.**
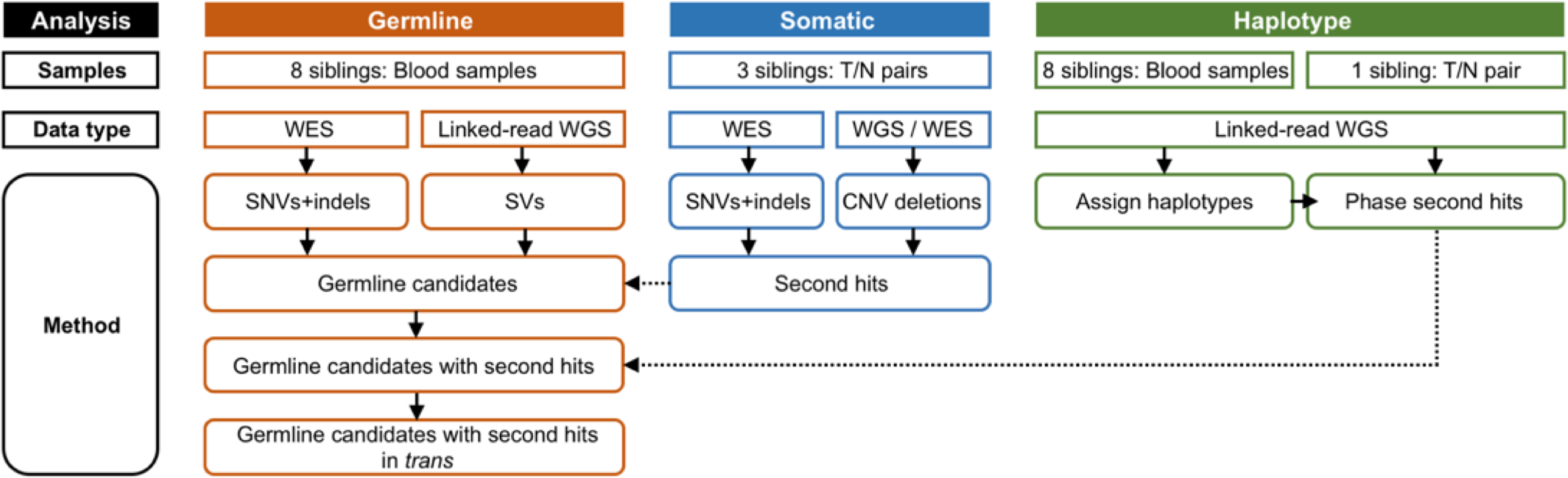
Overview of sequencing analysis methodology used to identify candidate germline variant in a family with CCA. Germline and somatic samples were sequenced with multiple sequencing technologies to detect variants. The resulting variants were integrated and filtered to identify a candidate causal mutation.

**Figure S2.**
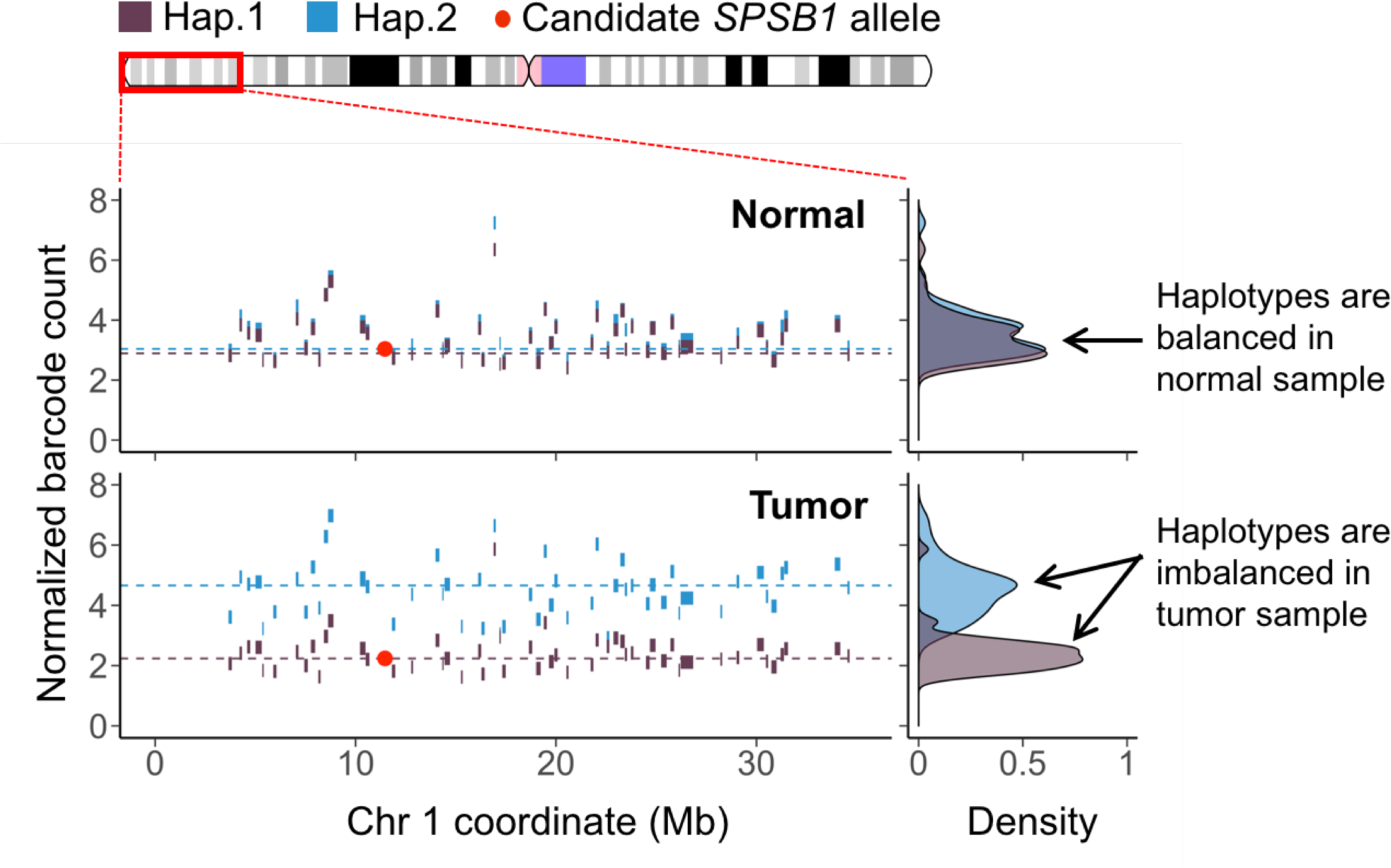
Extended haplotype of the 40 Mb deleted region of chromosome 1p in individual III:8. The blocks indicate the original fragmented haplotypes, and their color denotes their subsequent assignment to haplotypes covering many Mb. The candidate *SPSB1* allele exists in haplotype 1 (purple), which was the deleted haplotype in the tumor of this individual.

**Figure S3.**
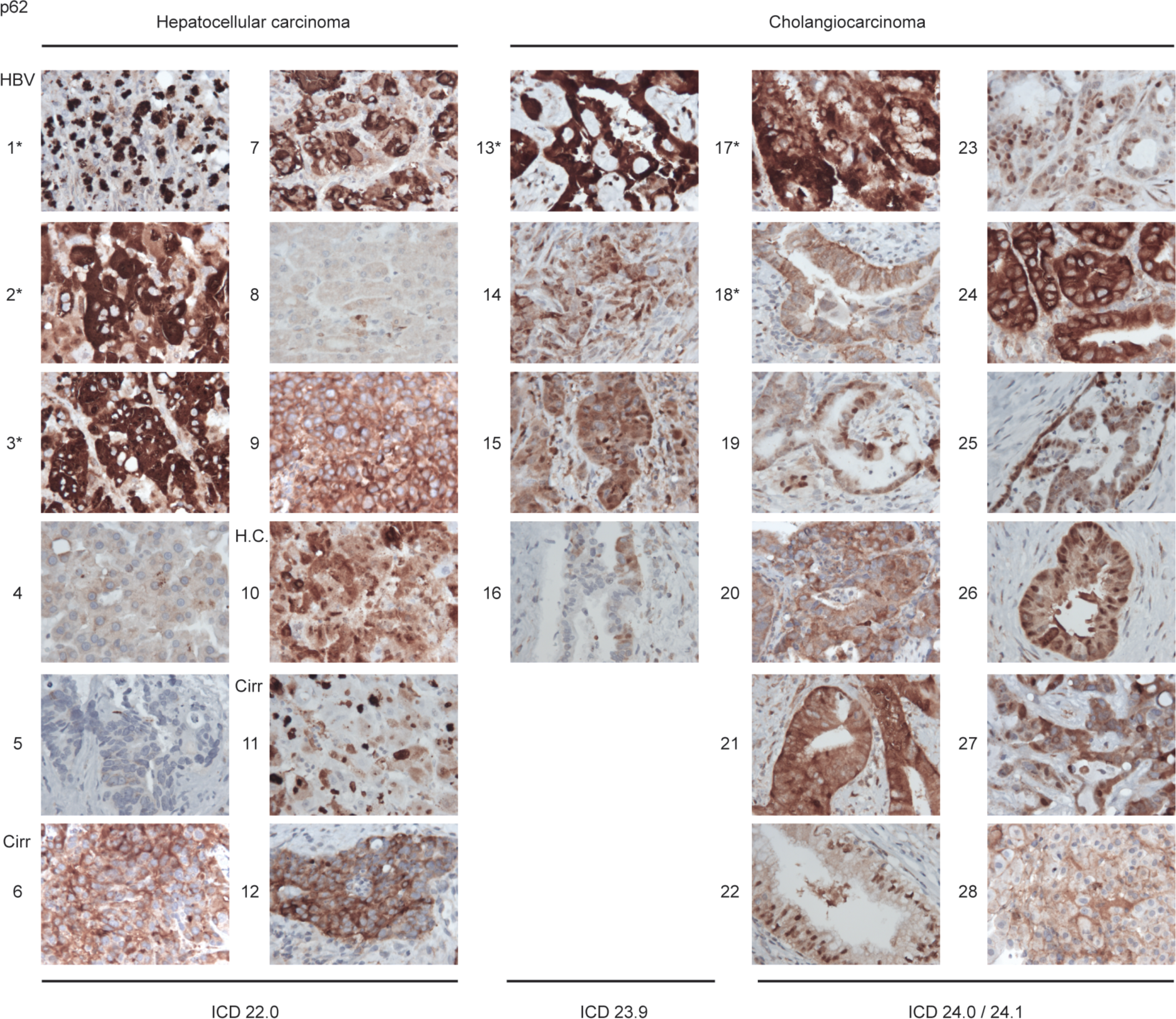
Increased p62 expression of HCC and CCA patients carrying the rs146589465-G variant. Sections from 28 paraffin-embedded HCC or CCA patient samples were stained with a p62 antibody. Numbers correlate with patient numbering in Table S20 and an asterisk indicates an rs146589465-G carrier. Information on liver disease is indicated; Cirr, cirrhosis; H.C., hemochromatosis; HBV, Hepatitis B virus. Samples are grouped by International Classification of Diseases (ICD) diagnoses information from the Icelandic Cancer Registry.

**Figure S4.**
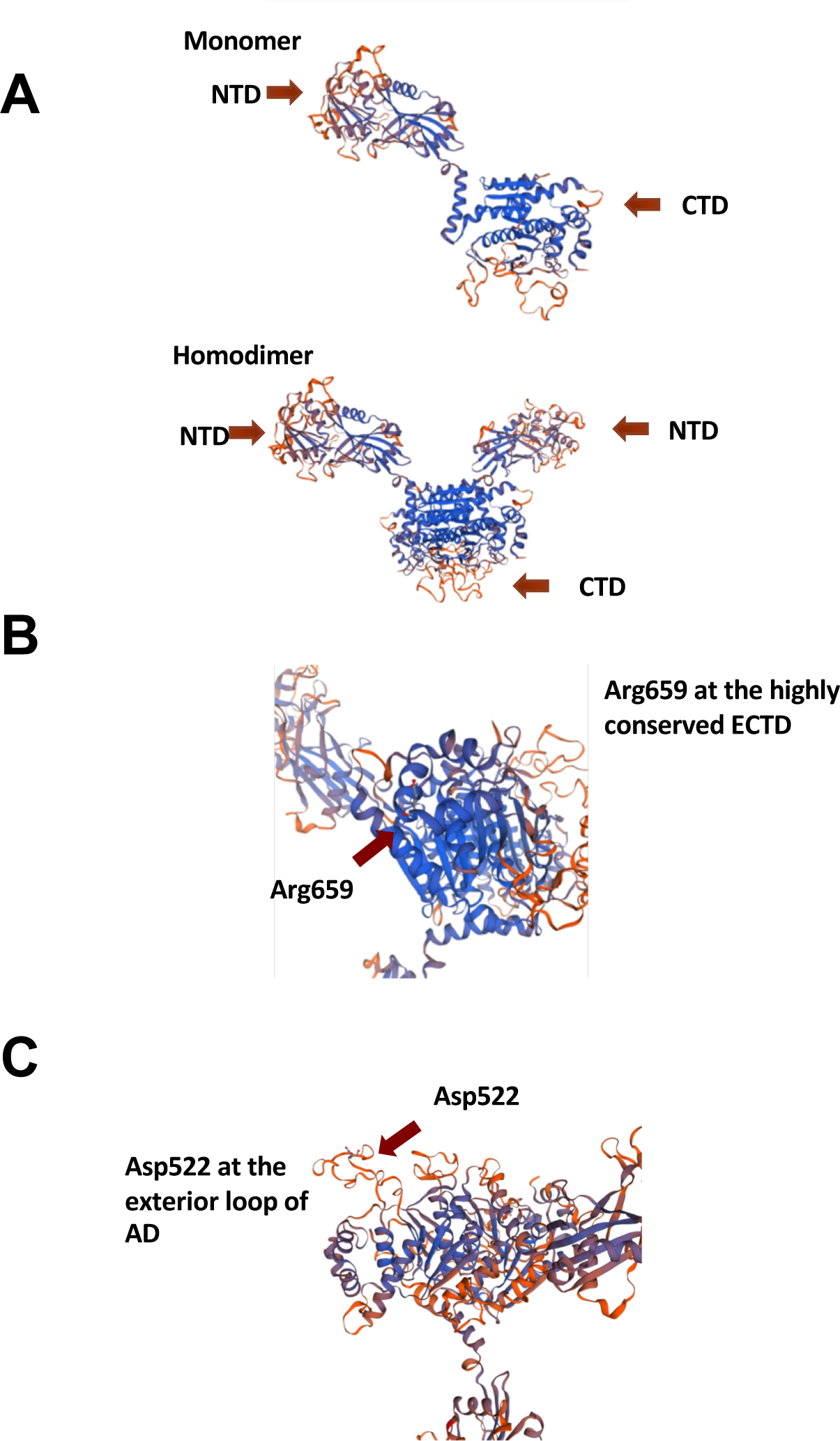
ATG7 protein structure and location of variants in the protein. (A) ATG7 functions as a homodimer. The locations of the CTD and NTD are indicated on the monomer and homodimer. **(B)** The Arg659 residue is located in the ECTD of ATG7. **(C)** The Asp522 residue is located in an exterior loop of the AD of ATG7.

**Figure S5.**
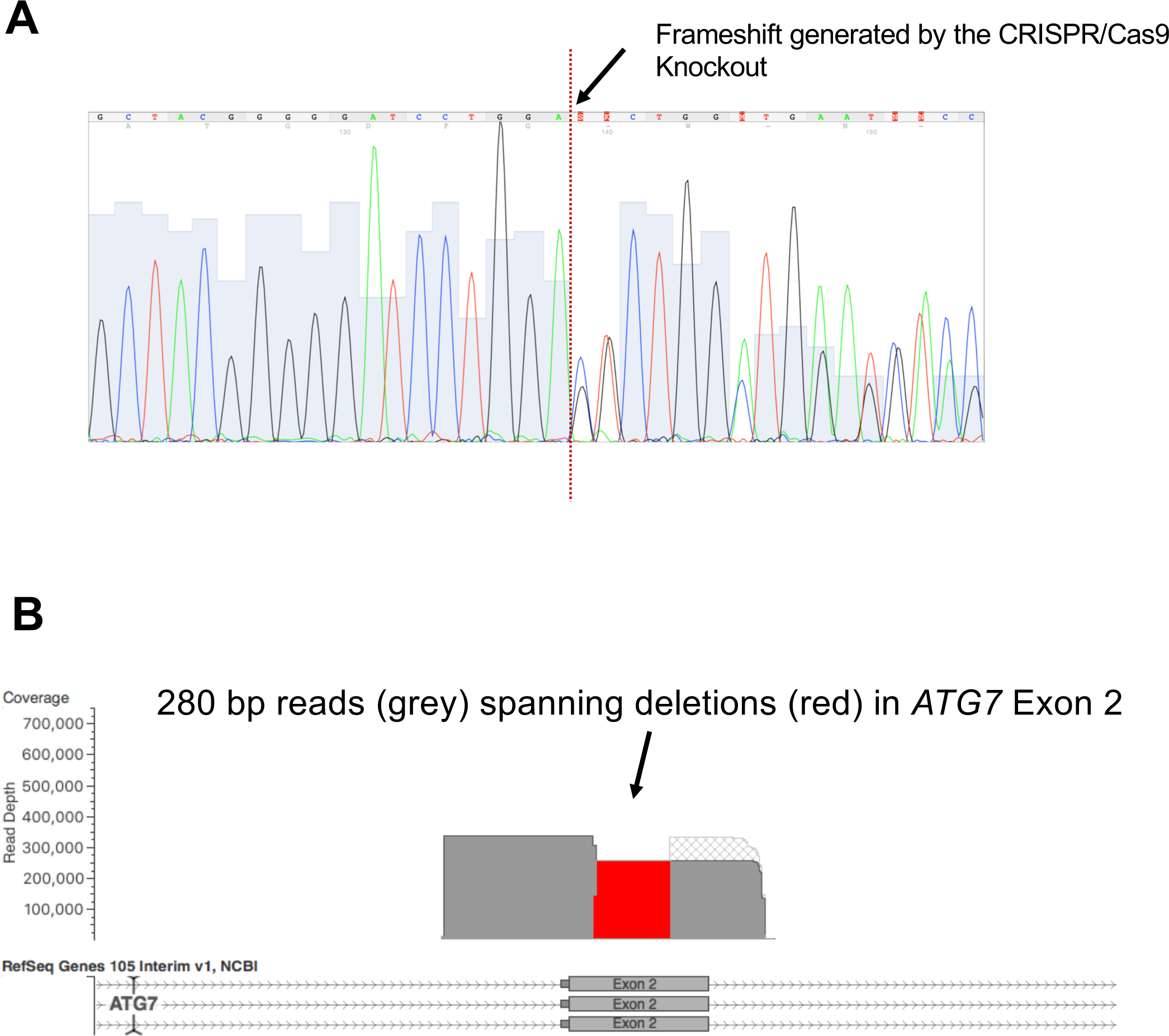
Confirmation of CRISPR deletion mutations in *ATG7* in the MMNK-1 cell line. **(A)** Sanger sequencing confirmed the presence of a deletion in an isogenic cell line. **(B)** Targeted amplicon sequencing with Illumina confirmed the presence of an 84-bp and 88-bp deletion in an isogenic MMNK-1 cell line (ATG7^-/-^).

**Figure S6.**
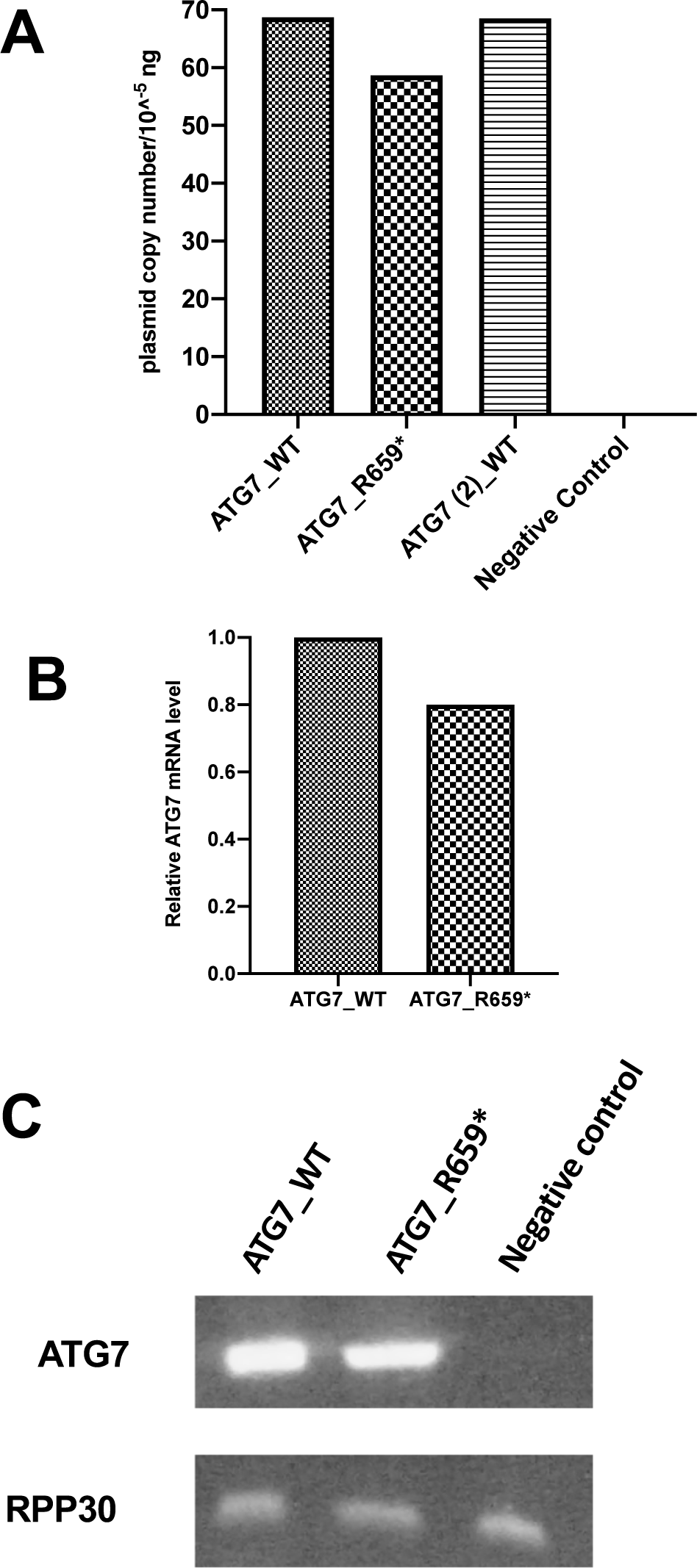
Quantification of plasmid input and mRNA expression level. **(A)** The copy number of each plasmid was quantified by ddPCR. The input amount of each plasmid was 10^-5^ ng/reaction. **(B)** Relative expression level of ATG7 mRNAs between WT and R659* expression in MMNK-1 ATG7^-/-^. The housekeeping gene *RPP30* was used for normalization. **(C)** ATG7 cDNA levels from MMNK-1 ATG7^-/-^ expressing WT and R659* plasmid using RT-PCR. Negative control is ATG7^-/-^ cells transfected with GFP expressing plasmids.

**Figure S7.**
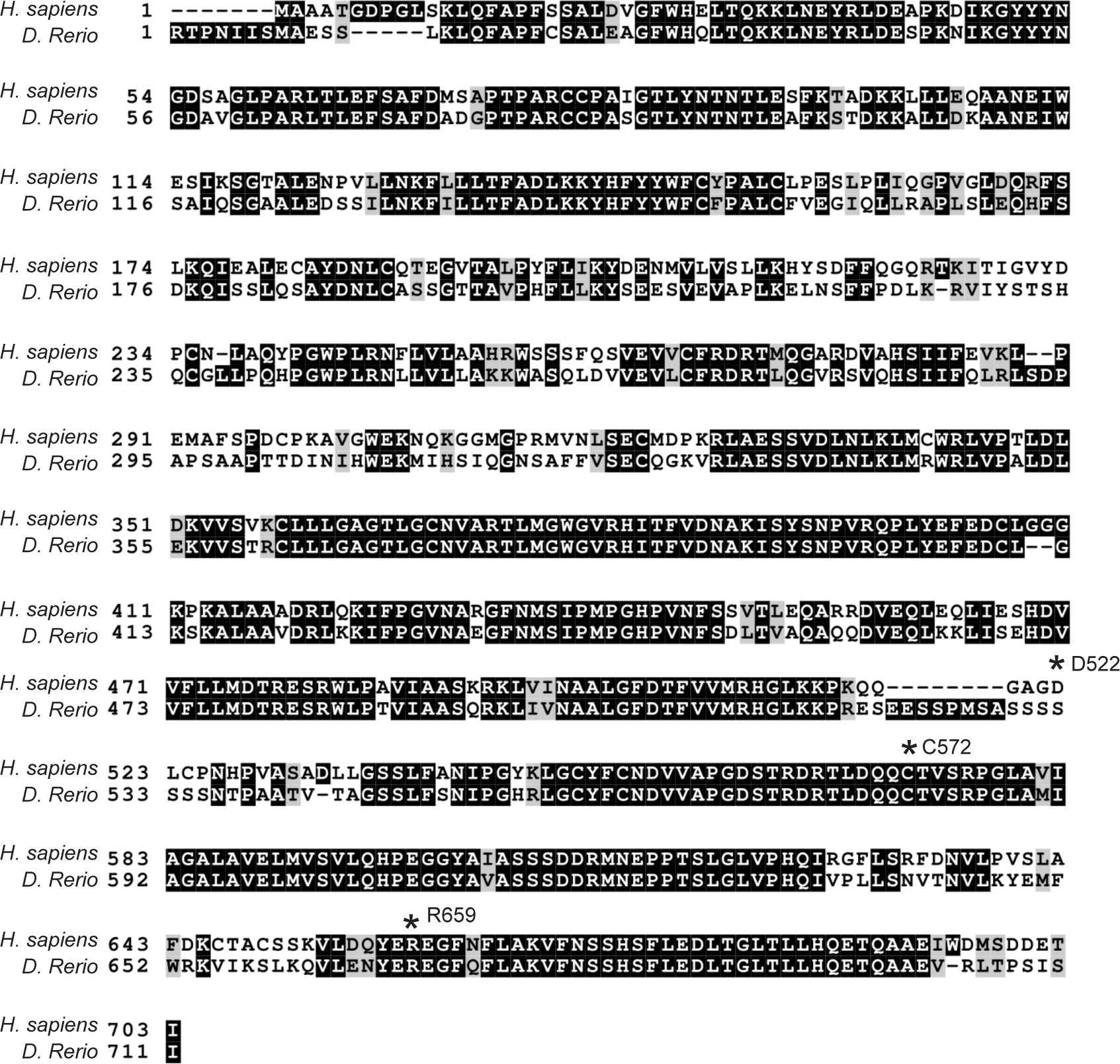
Sequence alignment of the ATG7 protein from human (*H. sapiens*) and zebrafish (*D. rerio*). Residues p.Asp522 (D522), p.Cys572 (C572) and p.Arg659 (R659) of human ATG7 are labeled with an asterisk. The alignment was performed with full length isoforms using EMBOSS Needle Pairwise Sequence Alignment^1^ and ExPASy BOXSHADE for visualizing the alignment.

